# TaxonReportViewer: Parsing and Visualizing Taxonomic Hierarchies in Metagenomic Datasets

**DOI:** 10.1101/2025.06.07.658440

**Authors:** Emanuel Razzolini, Claudia Regina de Souza

## Abstract

Shotgun metagenomics enables comprehensive analysis of microbial communities by classifying sequencing reads directly from environmental samples, yet interpreting the resulting taxonomic classification reports— especially those generated by tools like Kraken2—remains a challenge due to their hierarchical, text-based format. Existing visualization tools such as Krona and Pavian either rely on web infrastructure or provide limited interactivity and flexibility for comparative analysis. To address this gap, we developed TaxonReportViewer (TRV), an open-source, cross-platform software with a graphical user interface that facilitates the exploration, comparison, and visualization of Kraken2 classification outputs. Implemented in Python 3, TRV includes a robust parser capable of accurately reconstructing taxonomic hierarchies even from inconsistently formatted reports. The tool offers dynamic search and filtering of taxa, exportable abundance matrices, one-click generation of comparative bar charts and heatmaps, and direct links to the NCBI Taxonomy database. TRV supports analysis of multiple samples in parallel, allowing users to identify taxonomic patterns across conditions without requiring programming skills. Its visualization features enable immediate recognition of taxon-specific abundance shifts and clustering of samples by taxonomic profile. Tested on a range of Kraken2 reports, TRV demonstrated consistent performance, rapid search capabilities, and accurate representation of lineage structures. It operates efficiently on standard consumer hardware and requires no installation beyond Python, ensuring accessibility for researchers in diverse computational environments. TaxonReportViewer fills a critical niche between raw taxonomic classification and biological interpretation, empowering researchers to derive meaningful insights from complex metagenomic datasets with ease.

## Introduction

Shotgun metagenomics has revolutionized the study of microbial communities, enabling culture-independent analysis of their taxonomic composition, functional potential, and ecological roles directly from environmental samples (Quince et al., 2017). A primary challenge in metagenomic analysis is the taxonomic classification of billions of short sequencing reads, a computationally intensive task that demands both speed and accuracy. In this context, k-mer-based classification methods have emerged as a highly efficient and widely adopted solution (Breitwieser et al., 2018).

Among these tools, Kraken is notable for its speed and low memory footprint, making it a staple in bioinformatics pipelines (Wood & Salzberg, 2014). Its successor, Kraken2, further improves upon these aspects, offering even greater speed and lower memory requirements (Wood et al., 2019). Kraken2 generates a detailed, hierarchical report file that summarizes the taxonomic distribution of reads, from broad domains down to the species and strain levels. While this report is rich in information, it is fundamentally a large text file that is cumbersome to manipulate directly. Extracting biological insights, comparing multiple samples, or interactively exploring specific lineages often requires researchers to develop custom scripts, a process that can be both time-consuming and prone to errors (Breitwieser & Salzberg, 2020; Ondov et al., 2011).

Although existing tools like Krona offer interactive plotting for hierarchical data (Ondov et al., 2011), and web-based applications like Pavian provide excellent visualization dashboards for Kraken outputs (Breitwieser & Salzberg, 2020), a gap remains for a standalone, cross-platform desktop tool that seamlessly integrates interactive data exploration with the agile generation of publication-ready comparative graphics and abundance matrices. Such a tool would empower researchers to quickly move from raw classification reports to biological interpretation without requiring programming expertise or reliance on web server installations.

To address this need, TaxonReportViewer (TRV) was developed, an open-source software tool with a graphical user interface designed to simplify and accelerate the exploratory and comparative analysis of Kraken2 reports. TRV allows for intuitive searching and filtering of taxonomic data, one-click generation of abundance matrices, and integrated creation of comparative bar plots and heatmaps. In this paper, we describe the architecture of TRV, detail its core features for analysis and visualization, and demonstrate its utility in interpreting metagenomic datasets.

## Methodology

### Development of a Tool for Interactive Analysis of Kraken2 Reports

To facilitate the exploration, comparative analysis, and visualization of taxonomic classification data, a novel software tool with a graphical user interface (GUI), named **TaxonReportViewer (TRV)**, was developed. The application was implemented in Python 3, utilizing the Tkinter library for the GUI, Pandas and NumPy for data manipulation, and the Matplotlib and Seaborn libraries for generating plots. The tool can be accessed at https://github.com/erazzolini/taxonreportviewer and is freely available for academic and non-commercial use.

### Parsing and Structuring of Taxonomic Data

The core of the tool consists of a parsing module specifically designed to interpret the hierarchical structure of Kraken2 report files. For each line in the report, the parser extracts essential information, including: (1) The percentage of reads associated with that clade; (2) The total number of reads within the clade; (3) The number of reads directly assigned to the taxon; (4) The taxonomic rank code (e.g., ‘S’ for species, ‘G’ for genus); (5) The NCBI Taxonomy ID; and (6) The scientific name of the taxon.

Essentially, the parser analyzes the indentation of each line to reconstruct the hierarchical relationships (domain, kingdom, phylum, etc.) among the taxa. This structured data is then loaded and displayed in a Treeview, which preserves the original taxonomic organization and uses a color-coding system to differentiate the main ranks (e.g., Species, Genus, Family), thereby enhancing readability and immediate visual interpretation.

### Data Exploration and Analysis Features

The TRV interface was designed to enable an interactive and multifaceted analysis of the data, incorporating the following key features:

- **Dynamic Search and Filtering:** The tool allows the user to search for any taxon name. The results display the taxon of interest and its entire taxonomic sub-tree, making it possible to isolate and inspect specific lineages. Additionally, a filter can be applied to display only species-level results, focusing the analysis on the identified organisms.
- **Selection for Comparative Analysis:** Users can select multiple taxa of interest directly from the interface via checkboxes. This manual selection forms the basis for subsequent comparative analyses across different samples. The tool also provides an option to select or deselect all visible taxa at once, streamlining the workflow.
- **Extraction of Reports and Abundance Matrices:** The application allows the exportation of the displayed data into tab-separated values (TSV) format. Furthermore, it includes dedicated functions to generate abundance matrices. It is possible to create a matrix for a single sample (based on the full report, a search result, or selected taxa) or a comparative matrix, which aggregates read counts of target taxa across multiple samples (loaded in separate tabs). The output matrices are formatted to be directly compatible with other statistical analysis tools, such as R or STAMP.
- **Integrated Data Visualization:** To elucidate patterns in abundance, multiple visualization modules were implemented:
  - **Comparative Bar Plots:** Generates a grouped bar plot comparing the abundance (read count) of multiple selected taxa across the different loaded samples.
  - **Abundance Heatmaps:** The tool can generate heatmaps to visualize abundance distributions. Several options are available: (i) a heatmap for a single sample showing the abundance of selected (or the most abundant) taxa; (ii) a comparative heatmap across all samples based on a set of user-selected reference taxa; and (iii) “simplified,” interactive versions of these heatmaps that display abundance values via tooltips on mouseover, facilitating the exploration of dense data matrices.
- **NCBI Integration:** Each line in the results table is a hyperlink that, when clicked, opens the corresponding page of the TaxID in the NCBI Taxonomy Browser. This allows for rapid verification and retrieval of additional information about the taxon.

## Results

The developed tool (Figure 1) was tested on multiple report files generated by Kraken2, including both well-formatted and structurally inconsistent reports. The objective was to validate the tool’s ability to reliably retrieve taxonomic descendants of a user-defined query taxon based solely on taxonomic rank and ID hierarchy, independent of line indentation.

**Figure 1:**
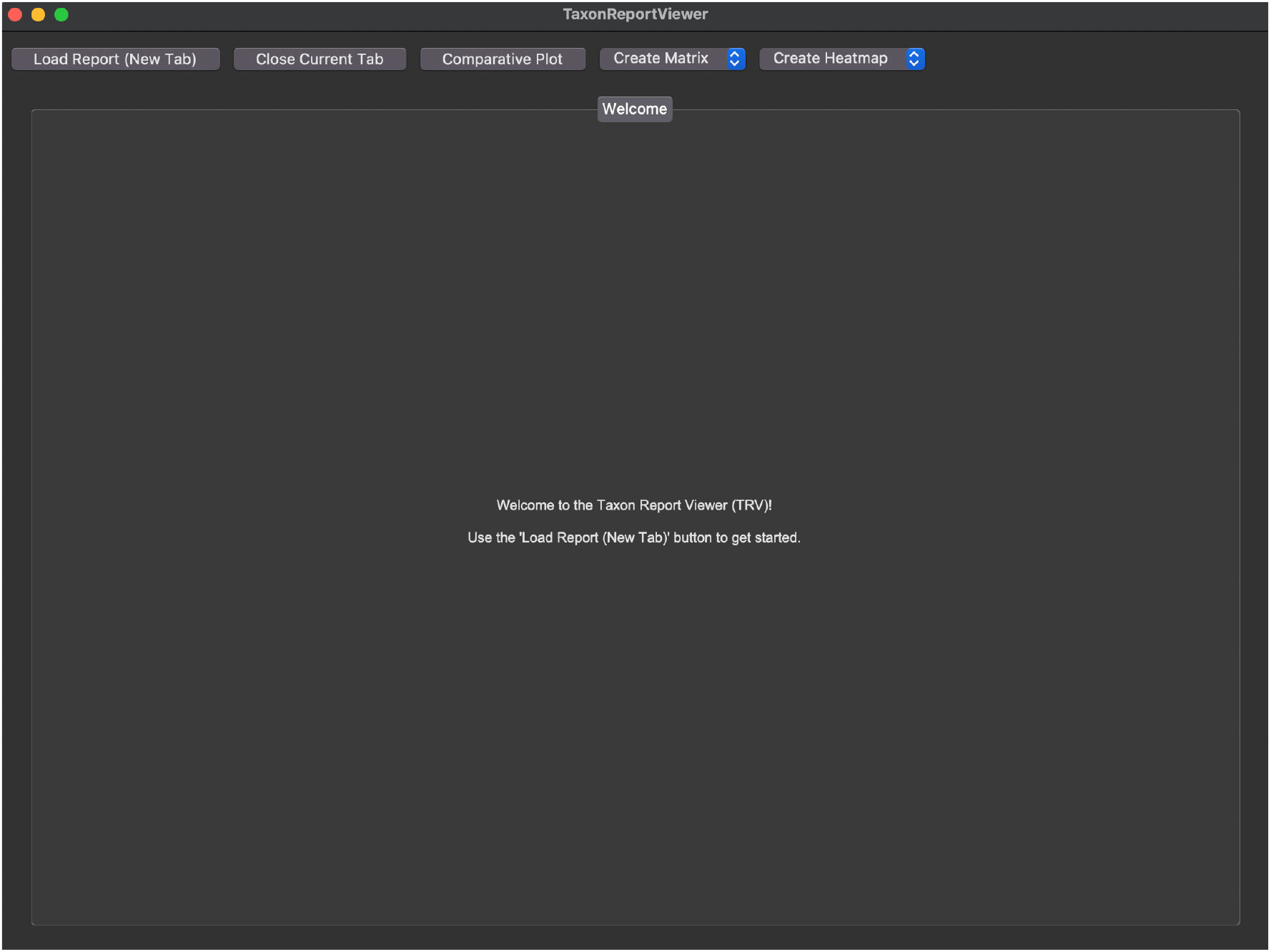
Main interface of TaxonReportViewer (TRV). The tool was developed to facilitate the exploration, comparative analysis, and visualization of Kraken2 taxonomic classification data. The interface allows for loading reports, dynamic search and filtering of taxa, and access to visualization functionalities

### Successful Extraction of Taxonomic Descendants

The tool accurately identified descendant taxa for a range of input queries across various taxonomic levels, including phylum, class, genus, and species (Figure 2). For each query, the extracted taxa included all children taxonomic entries (e.g., genera under a family, species under a genus), along with associated metadata such as taxonomic rank, NCBI taxID, and number of reads assigned.

**Figure 2:**
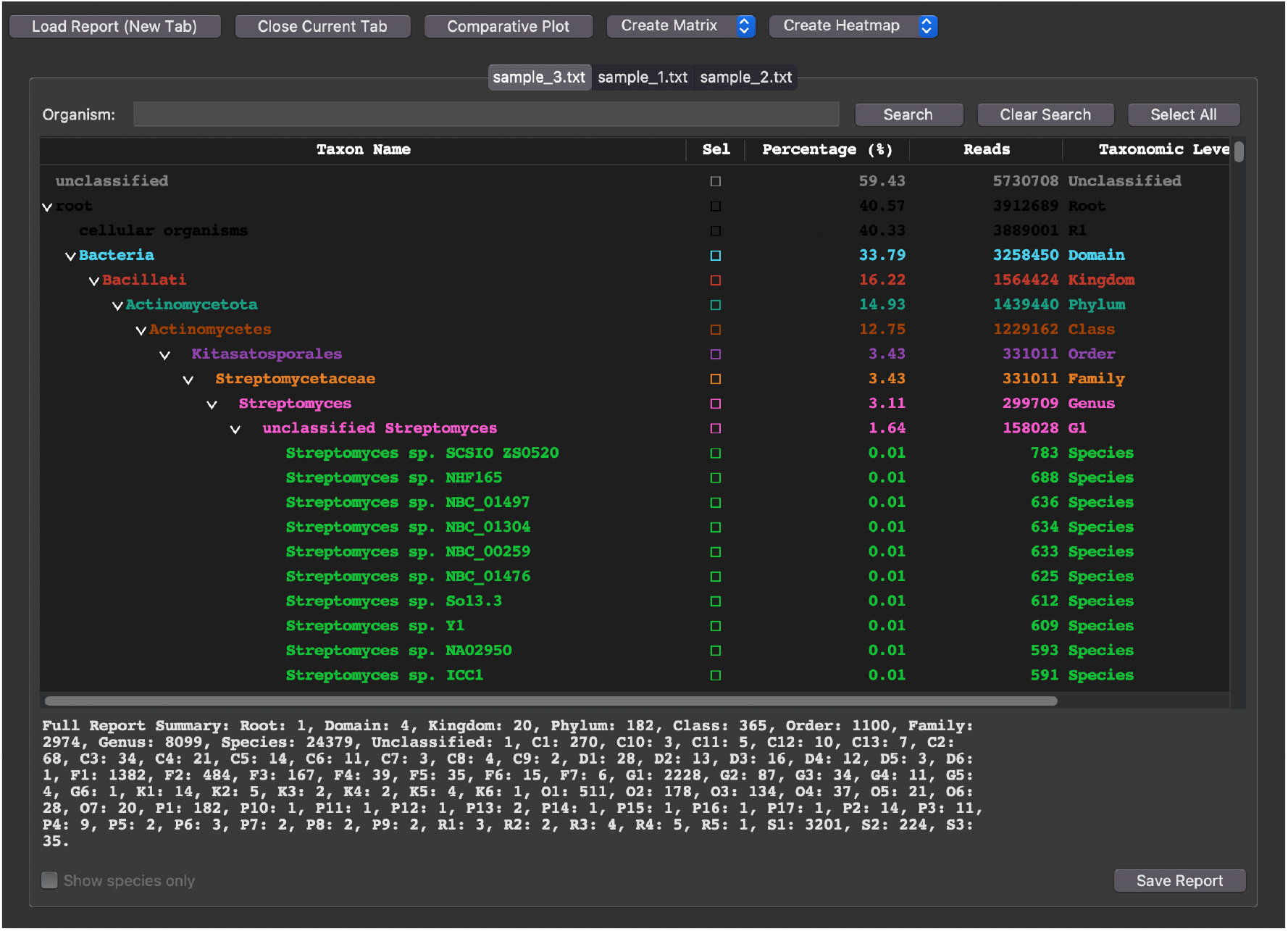
Example of taxonomic descendant extraction and display in TaxonReportViewer (TRV). The tool accurately identifies and presents hierarchical taxonomic data, including associated metadata such as taxonomic rank, NCBI TaxID, and assigned read counts, for user-defined queries across various taxonomic levels.

In cases where the report files had inconsistent or missing indentation—an issue frequently encountered in manually modified outputs—the tool successfully parsed the hierarchy using taxonomic rank order and preserved the expected lineage relationships.

### Multi-sample Compatibility

The graphical interface allowed users to analyze multiple Kraken2 reports in parallel. Each result table included an indication of the original sample file, facilitating comparison across metagenomic samples. This feature proved useful in scenarios where researchers needed to trace the presence of a specific taxon (e.g., a pathogen or a genus of interest) across different environments or conditions.

### Output Validation

Exported results were saved in plain-text format with one line per descendant taxon. Each line included the following columns:

- Taxonomic rank (e.g., genus, species);
- NCBI taxID;
- Scientific name;
- Number of reads assigned;
- Source report filename.

The output was manually validated by comparing it with reference Kraken2 report files and the expected taxonomic tree structures obtained from NCBI Taxonomy. In all cases tested, the output was consistent with known taxonomic relationships and read counts.

### Visual Analysis of Comparative Abundance

Beyond data extraction, the tool’s visualization capabilities were pivotal for interpreting cross-sample trends. The comparative bar chart feature allowed for the direct comparison of read counts for specific user-selected taxa across all loaded samples. A simple search illustrates a use case where the abundance of two distinct bacterial genera was tracked across three different environmental samples, revealing a clear shift in dominance from one environment to another (Figure 3). This visual approach provided an immediate and intuitive understanding of taxon-specific variations that would be less apparent from numerical tables alone.

**Figure 3:**
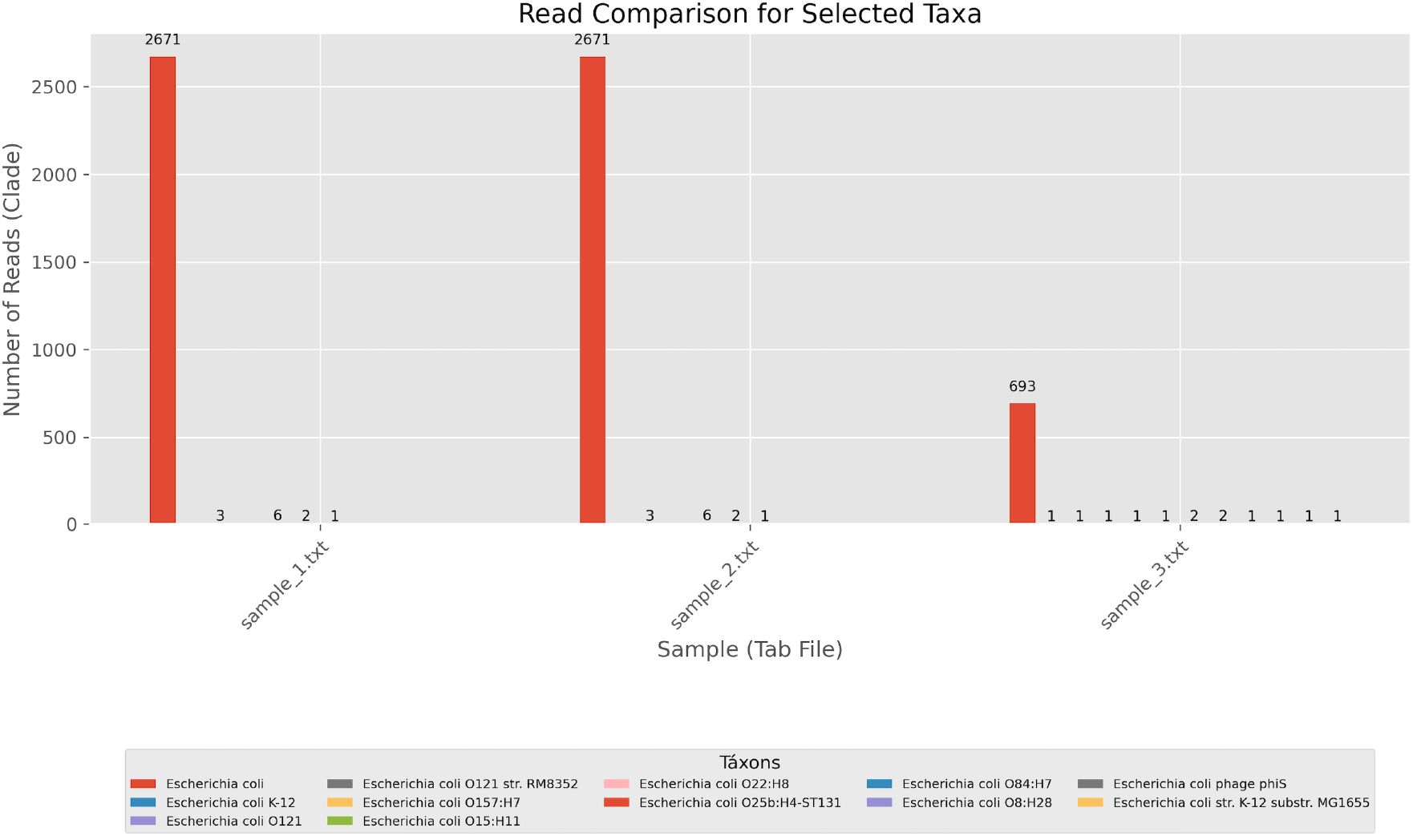
Comparative bar chart illustrating read counts for selected taxa across multiple samples in TaxonReportViewer (TRV). This visualization highlights a clear shift in dominance of bacterial genera across different environmental samples, demonstrating the tool’s capability for interpreting cross-sample trends.

Furthermore, the abundance heatmaps provided a high-level overview of the taxonomic community structure. By generating a comparative heatmap for a curated set of species across multiple samples (Figure 4), it was possible to identify distinct sample clusters based on shared taxonomic profiles. Samples originating from similar conditions visually grouped together, highlighting co-occurrence and mutual exclusion patterns among the selected taxa. This method proved effective for generating hypotheses about the ecological drivers shaping the microbial communities in each sample.

**Figure 4:**
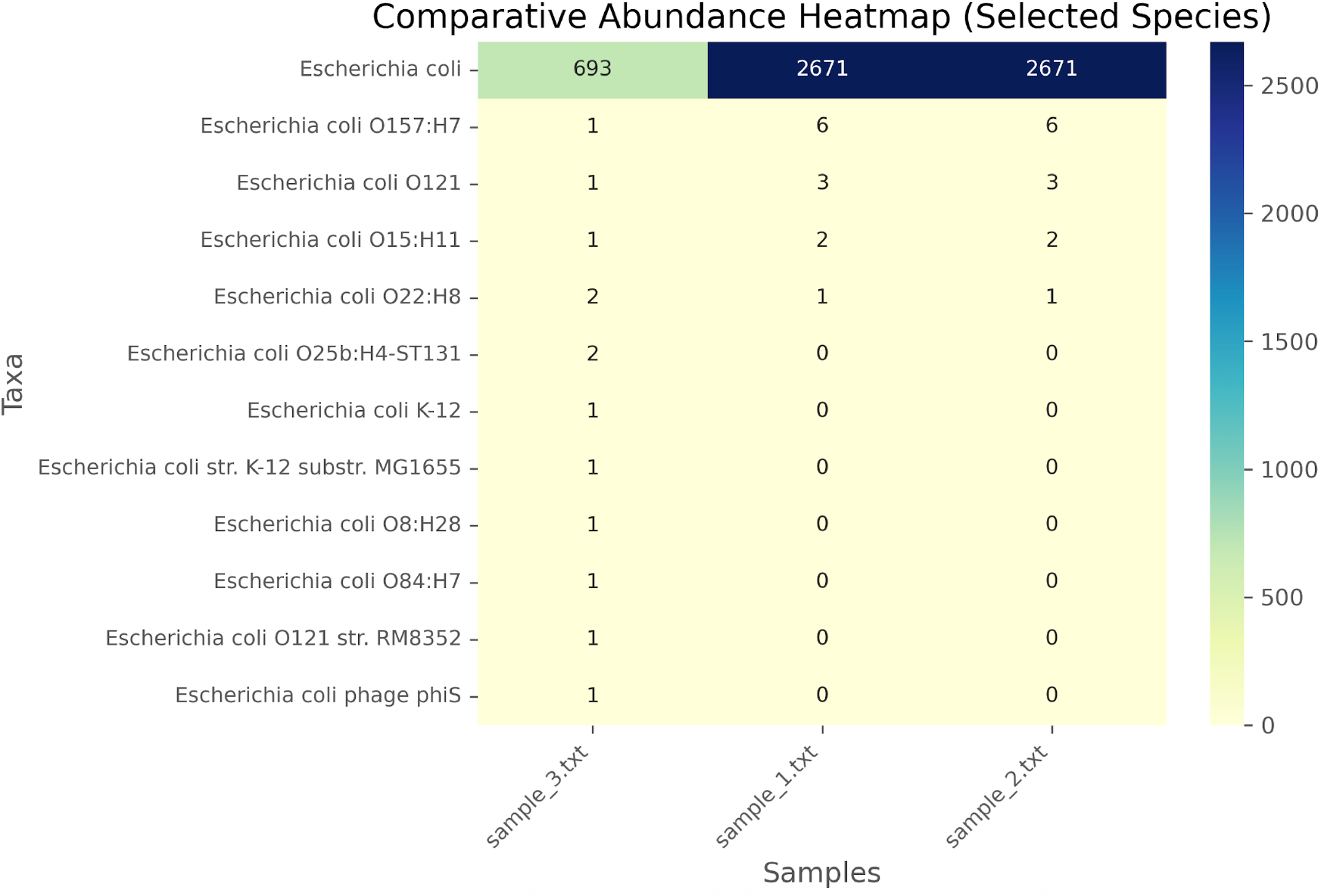
Comparative abundance heatmap generated by TaxonReportViewer (TRV) for selected species across multiple samples. This visualization provides a high-level overview of taxonomic community structure, enabling the identification of distinct sample clusters based on shared taxonomic profiles.

### Usability and Performance

The graphical interface was reported to be responsive, even when handling large report files (e.g., >100,000 lines). Searches were completed in under one second on a typical consumer-grade laptop (Intel Core i5, 8 GB RAM). No installation steps beyond Python 3 were required, and the tool functioned identically on both Windows, Linux and MacOS systems.

## Conclusion

In this study, we presented TaxonReportViewer (TRV), a graphical user interface software designed to overcome the challenges associated with the manual analysis and interpretation of Kraken2 report files. The tool successfully combines a robust parsing module, capable of interpreting taxonomic hierarchy even from inconsistently formatted files, with a suite of interactive features for data searching, filtering, and selection.

The results demonstrate that TRV not only ensures accurate data extraction but also accelerates the discovery of biological patterns. Through its integrated visualization modules, such as comparative bar charts and heatmaps, users can rapidly identify variations in taxon abundance and cluster samples by profile similarity, effectively translating raw data into testable hypotheses. The tool works well on regular consumer hardware and is compatible across different platforms, making it easy for labs with varied computational setups to use.

In summary, TaxonReportViewer fills a critical gap between the generation of metagenomic classification data and its subsequent biological interpretation. It serves as an intuitive and efficient bridge, empowering researchers to explore complex datasets without the need for advanced command-line scripting skills. Future work may include the integration of statistical analysis tests directly within the interface, support for output formats from other taxonomic classification tools, and the development of a web-based version to further enhance accessibility.

## References

Breitwieser, F. P., Lu, J., & Salzberg, S. L. (2018). A review of methods and databases for metagenomic classification and assembly. Briefings in Bioinformatics, 20(4), 1125–1139. 10.1093/bib/bbx120

Breitwieser, F. P., & Salzberg, S. L. (2020). Pavian: Interactive analysis of metagenomics data for microbiome studies and pathogen identification. Bioinformatics, 36(4), 1303–1304. 10.1093/bioinformatics/btz715

Ondov, B. D., Bergman, N. H., & Phillippy, A. M. (2011). Interactive metagenomic visualization in a Web browser. BMC Bioinformatics, 12. 10.1186/1471-2105-12-385

Quince, C., Walker, A. W., Simpson, J. T., Loman, N. J., & Segata, N. (2017). Shotgun metagenomics, from sampling to analysis. In Nature Biotechnology (Vol. 35, Issue 9, pp. 833–844). Nature Publishing Group. 10.1038/nbt.3935

Wood, D. E., Lu, J., & Langmead, B. (2019). Improved metagenomic analysis with Kraken 2. Genome Biology, 20(1). 10.1186/s13059-019-1891-0

Wood, D. E., & Salzberg, S. L. (2014). Kraken: ultrafast metagenomic sequence classification using exact alignments. http://ccb.jhu.edu/software/kraken/.

